# Rapid evolution of gained essential developmental functions of a young gene via interactions with other essential genes

**DOI:** 10.1101/226936

**Authors:** Yuh Chwen G. Lee, Iuri M. Ventura, Gavin R. Rice, Don-Yuan Chen, Manyuan Long

## Abstract

New genes originated relatively recently and are only present in a subset of species in a phylogeny. Accumulated evidence suggests that new genes, like old genes that are conserved across species, can also take on important functions and be essential for the survival and reproductive success of organisms. While there are detailed analyses of the mechanisms underlying gained fertility functions by new genes, how new genes rapidly became essential for viability remains unclear. We focused on a young retro-duplicated gene (*CG7804*, which we named *Cocoon*) in *Drosophila* that originated three million years ago. We found that, unlike its evolutionarily conserved and broadly expressed parental gene, *Cocoon* has evolved rapidly under positive selection since its birth and accumulates many amino acid divergences at functional sites from the parental gene. Despite its young age, *Cocoon* is essential for the survival of *D. melanogaster* at multiple developmental stages, including the critical embryonic stage, and its expression is essential in different tissues from its parental gene. Functional genomic analyses found that *Cocoon* gained multiple DNA binding targets, which regulates the expression of genes that have other essential functions and/or have multiple gene-gene interactions. Our observations suggest that *Cocoon* acquired essential function to survival through forming interactions that have large impacts on the gene interaction network. Our study is an important step towards deciphering the evolutionary trajectory by which new genes functionally diverge from the parental gene and become essential.

## Introduction

Every gene in an organism’s genome must have arisen at some time point in the past. The origination of new genes is an important evolutionary process contributing to the dynamic turnover of genes in genomes over the phylogeny. This dynamic gene turn over has been widely documented in *Drosophila* (Zhang *et al.* 2010), primates (Demuth *et al.* 2006; Zhang *et al.* 2011), and plants (Moore and Purugganan 2005) (reviewed in (Kaessmann 2010). Because of their recent origin, new genes are only present in a subset of species in a phylogeny and a naïve expectation would predict they have dispensable functions and are not essential to an organism’s fitness (e.g. (Ashburner *et al.* 1999)). However, evidence accumulated from a variety of eukaryotic species shows that new genes can quickly become essential for an organism’s viability and fertility (Chen *et al.* 2010; Cooper and Kehrer-Sawatzki 2011; Ding *et al.* 2012; Ranz and Parsch 2012; Charrier *et al.* 2012; Dennis *et al.* 2012; Ross *et al.* 2013; Reinhardt *et al.* 2013; Gubala *et al.* 2017).

One of the mechanisms by which new genes arise is through duplication, in which a copy of a gene is created through either DNA or RNA intermediates in the genome. Many evolutionary fates have been predicted for the duplicated (new) and original (parental) genes, grossly pseudo-functionalization, neo-functionalization, or sub-functionalization (Ohno 1970; Lynch and Conery 2000; Innan and Kondrashov 2010). Despite the convenient conceptual distinction, it is often challenging to distinguish between these alternative models due to the fact that the past evolutionary trajectories are usually unknown or hard to decipher. Several in-depth analyses of the evolutionary steps leading to gained novel fertility functions of duplicated genes (Loppin *et al.* 2005; Heinen *et al.* 2009; Ding *et al.* 2010; Yeh *et al.* 2012; Chen *et al.* 2012) have shed light on the initial evolutionary processes of gained essential function supported by new genes. In contrast, few studies have focused on viability (e.g. (Ross *et al.* 2013)). Many genes responsible for essential viability functions (e.g. development of body plan in *Drosophila* embryos (Stauber *et al.* 1999)) are identified as ancient gene duplicates (reviewed in (Chen *et al.* 2013)), suggesting new genes indeed can gain a critical role in the most essential and core functions of organisms. Yet, the past evolutionary trajectories of gained essential viability function by new genes, and whether that is similar to those of essential fertility function, still needs further investigation.

A potential mechanism by which duplicated genes become essential is by integrating into the cellular genetic network, through either new protein-protein or new protein-nucleic acids interactions with pre-existing genes. Indeed, new genes with essential fertility functions were discovered to locally or globally reshaped the regulatory network (Matsuno *et al.* 2009; Ding *et al.* 2010; Chen *et al.* 2012). A genome-wide study found that many young duplicated genes that are essential for survival also acquired multiple novel protein-protein interactions (Chen *et al.* 2010), which is consistent with observations that genes with many interaction partners (hub genes) are more likely to have essential functions ((Jeong *et al.* 2001; Yu *et al.* 2004; Batada *et al.* 2007; Blomen *et al.* 2015) and reviewed in (Barabási and Oltvai 2004; Barabási *et al.* 2011)). Paradoxically, comparisons of ancient orthologous genes report that the accumulation of gene-gene interactions is a slow evolutionary process (Kim *et al.* 2012). This raises the question of how, in a short evolutionary time, new genes can acquire multiple new interactions, and how this leads to their essential functions.

In this study, we characterized the evolutionary history and function of a young duplicated gene that quickly became essential for the *survival* of *Drosophila melanogaster.* This young gene (*CG7804*) duplicated from another essential gene (*TBPH*, also known as *TDP-43 human homolog* or CG10327) through retrotransposition less than four million years ago (Zhang *et al.* 2010), and is present in few *Drosophila* species. The especially young age of *CG7804* offers a rare opportunity to investigate the initial evolutionary steps of gaining essentiality by new genes. The parental gene, *TBPH*, is highly conserved among animals (Ayala *et al.* 2005; Li *et al.* 2010), its null mutant was found lethal in *Drosophila* (Feiguin *et al.* 2009; Lin *et al.* 2011; Hazelett *et al.* 2012), and *TBPH* mutant in human is associated with neuronal diseases (Sreedharan *et al.* 2008). *TBPH* is shown to bind to nucleic acid (Kuo *et al.* 2009), influencing the splicing (Buratti and Baralle 2001; Ayala *et al.* 2006; Bose *et al.* 2008) and transcriptional regulation (Ayala *et al.* 2008) of many genes. On the other hand, little is known about the duplicated gene, *CG7804.* We found that unlike its evolutionarily conserved, broadly expressed parental gene, *CG7804* has evolved rapidly under positive selection since its birth. Despite its young age, functional analyses show that *CG7804* is essential for the survival of *D. melanogaster* at multiple developmental stages, including the critical embryonic stage. In particular, its expression is essential in different tissues from its parental gene. RNA-seq and ChIP-seq analysis suggests that *CG7804* acquired essential function to survival through gaining DNA binding targets that influence the expression of genes that have other essential function (i.e. mutant lethal) as well as involve in a large number of protein-protein/gene-gene interactions. Our study is an important step towards deciphering the evolutionary trajectory by which duplicated genes functionally diverge from the parental gene and become essential.

## Results

### *CG7804* evolved quickly since its origination

*CG7804* (on chr3L) originated 1.5 to 3.5 million years ago via an RNA intermediate (retrotransposition) from *TBPH* (on chr2R). Among the sequenced 12 *Drosophila* species (Clark *et al.* 2007), is only present in *D. melanogaster, D. simulans*, and *D sechellia* (Zhang *et al.* 2010) (Figure 1A). Despite its short evolutionary history, we detected a burst of 100 amino acid substitutions during the initial one million years after the origination of *CG7804* by using maximum-likelihood method (Yang 2007) (Figure 1B). A likelihood ratio test found that the *dN/dS* ratio is significantly different between the clade of *CG7804* and the clade of *TBPH* (likelihood-ratio test, *p-value* = 0.003; Figure 1B). The *dN/dS* ratio of the *CG7804* clade (2.88) is much greater than one, which is usually interpreted as a signature of positive selection. In contrast, *dN/dS* ratio of the *TBPH* clade (0.041) is much smaller than one, consistent with purifying selection constraining its evolution. Furthermore, we found a significant excess of amino acid substitutions in *CG7804* using the McDonald-Kreitman test ((McDonald and Kreitman 1991); *Fisher’s Exact Test*, *p-value* = 0.0011) and 75.8% of these amino acid substitutions are inferred to have been fixed by positive selection (*α*, (Smith and Eyre-Walker 2002)), suggesting *CG7804* has been under positive selection. On the contrary, no evidence supports that *TBPH* is under positive selection (McDonald-Kreitman test, *p-value* = 0.27).

**Figure 1.**
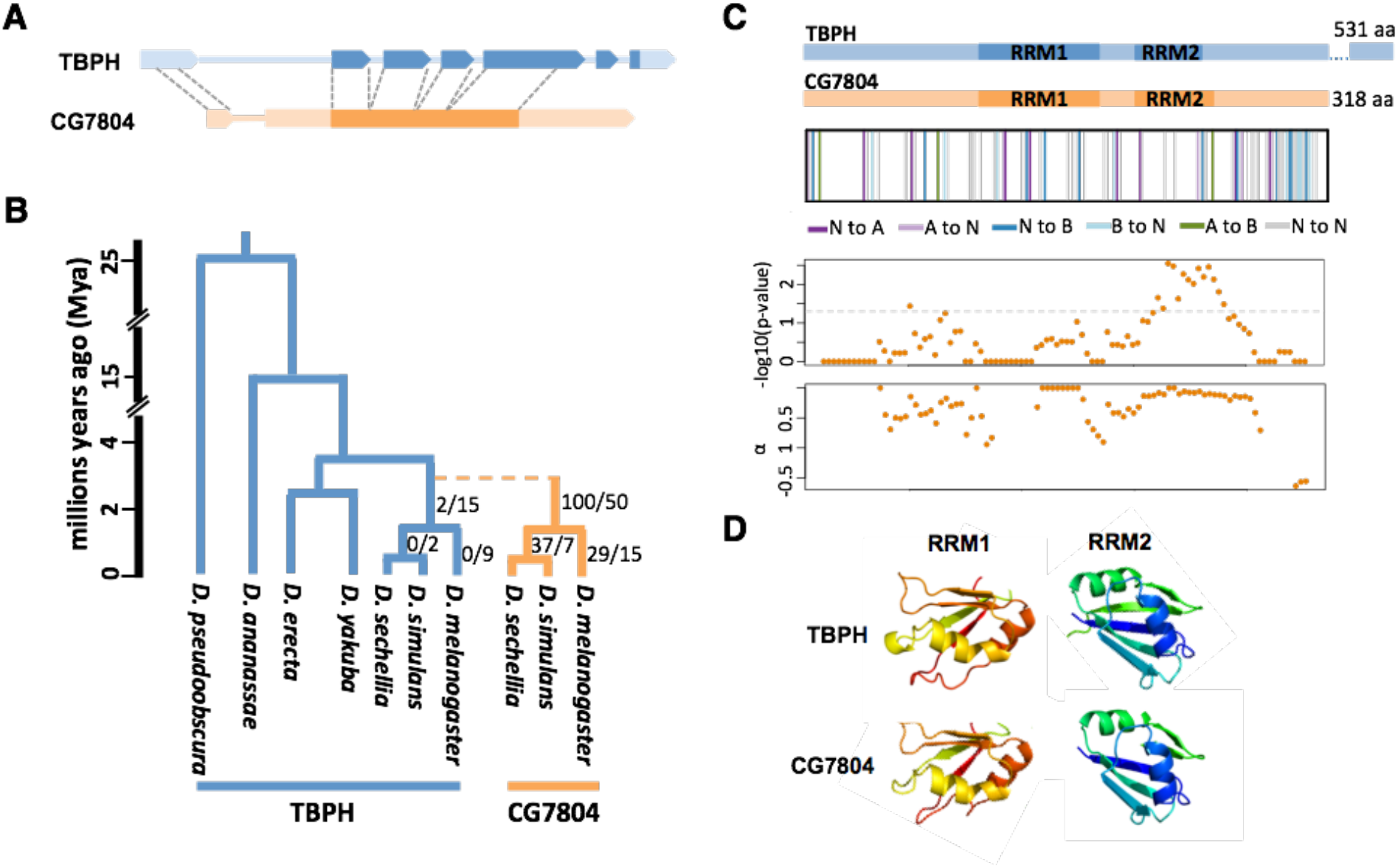
Structural and evolutionary history of *CG7804* and *TBPH*. (A) Exon-intron structure of *TBPH* (blue) and *CG7804* (orange). Filled boxes represent exons while lines represent introns. Because *CG7804* originated through a retrotransposition event, it lacks most of the introns of *TBPH* and some of its noncoding sequences do not share homology with *TBPH.* (B) The duplication event of *CG7804* from *TBPH* is denoted as dashed line in the phylogeny. The clades of *CG7804* and *TBPH* are in orange and blue respectively. The number of amino acid substitutions to number of synonymous substitutions inferred by PAML (see text) is denoted at right to branches. The dating of the species phylogeny is from (Obbard *et al.* 2012). (C) The structures of the *amino acid* sequences of *TBPH* and *CG7804* are shown at the upper panel. In the second panel, the divergent amino acids of *CG7804* from *TBPH* are denoted as vertical lines. Different colors are different changes in amino acid chemical properties (N: non-charged, A: acidic, B: basic). The third and fourth panels show the results of sliding window *MK* test of *CG7804* (window size 99 bp, step 9 bp), including -log10 *p-value* of the *MK test* and the estimated proportion of amino acid fixations driven by positive selection (α). Note that the coordinates of four panels are aligned. (D) Predicted structures of the RRMs of *TBPH* and *CG7804* are shown. The 3D structures of the first (RRM1) and the second (RRM2) RNA-recognition motifs were predicted using Phyre (Kelley *et al.* 2015). The rainbow color is from N (red) to C (blue) termini.

The fast evolution of *CG7804* since its origination resulted in the high divergence in amino acid sequences between it and *TBPH* (28.6%, 91 out of 318 amino acids). Many of these amino acid substitutions lead to changes in protein charges (Figure 1C). As a comparison, *TBPH* paralogs from *D. melanogaster* and *D. yakuba*, a species in which *CG7804* is absent, diverge only 5.0% in amino acid sequences (27 amino acids). Furthermore, *TBPH* in *D. melanogaster* accumulated only 2 amino acid substitutions since the duplication event of *CG7804* (Figure 1B).

*TBPH* has two RRM (RNA recognition motif) domains, which has been demonstrated to bind to both RNA and DNA (Kuo *et al.* 2009). *CG7804* is predicted to possess the same nucleic acid binding domains (amino acid 109-174, 194-239) with similar overall structures (Figure 1D). Interestingly, sliding window MK test analysis found that regions with significant MK results and/or a large proportion of amino acid substitutions fixed by positive selection overlap with either of the two predicted RRM domains of *CG7804* (Figure 1C). Furthermore, at least two amino acid differences between *CG7804* and *TBPH* are likely to have substantial functional effects. Lukavsky et al. (Lukavsky *et al.* 2013) experimentally identified that, in human *TBPH*, Met132 and the salt bridge between the residues Arg151 and Asp247 are important for nucleic acid-binding affinity. Both of these residues are highly conserved among *TBPH* orthologs in animals (Supplementary Figure 1). However, in *CG7804*, these key residues were replaced by amino acids with different charges (polar Met132 was substituted with positively charged lysine, negatively charged Asp247 was substituted with the positively charged lysine, Supplementary Figure 1), both likely led to diverged nucleic acid binding targets and/or functional role of *CG7804* from *TBPH.* Taken together, these results suggest that, whereas the parental gene remained highly constrained at the amino acid sequences, *CG7804* quickly accumulated many amino acid substitutions, even at functionally important domains and sites.

### *CG7804* is essential for the survival of *D. melanogaster*

Because our evolutionary genetic analysis supports positive selection acting on *CG7804*, we predicted that it gained function since its origination, instead of being pseudogenized or retained simply due to selection for increased dosage of expression, in which case selection will be mainly acting on the regulatory sequences. We employed GAL4/UAS system, and first used ubiquitous GAL4 drivers (Act5C-GAL4 and Tub-GAL4) to knock down the expression of *CG7804* and *TBPH* individually (see Materials and Methods). Consistent with previous study using *TBPH* knockout flies (Feiguin *et al.* 2009; Lin *et al.* 2011; Hazelett *et al.* 2012), *TBPH* knockdown found the gene being essential for the survival of *D. melanogaster* (lethality rate: 97.5% with Act5C-GAL4 and 86.6% with Tub-GAL4, see Materials and Methods). Surprisingly, despite originating recently on an evolutionary timescale, expression knockdown of *CG7804* also led to very low survival rate (lethality rate: 94.7% with Act5C-GAL4 and 95.5% with Tub-GAL4). For *CG7804* knockdown, most of the lethality happens at the stage from larvae to adults (Figure 2A). Indeed, we found that flies could not develop pass the pharate adult stage and identified many eclosion lethal incidences (i.e. flies could not emerge and were found stuck and dead half way in pupal cases, Figure 2B). Many other flies that eclosed were dead in *Drosophila* culture media. A detailed tracking of pupa identified that, while 91.9% of the wildtype pupa successfully eclosed and survived, only 17.7% of the *CG7804* knockdown individuals who reached the pupal stage did so (Figure 2C). To test if the high lethality associated with *CG7804* knockdown is caused by flies unable to first open the pupal cases, we manually removed the pupal cap for both *CG7804-knock* down and wildtype flies. Manual removal of pupal cap led to a slight increase in pupal lethality for wildtype flies (increased from 5% to 12.5%; Figure 2C). On the other hand, pupal cap removal decreased the lethality rate of *CG7804-*knockdown pupa from 76.6% to 60% (Figure 2C). Yet, even after considering the increased lethality due to pupal cap removal (~7.5% in wildtype), 52.5% *CG7804* knockdown pupa still did not reach adulthood (Figure 2C).

**Figure 2.**
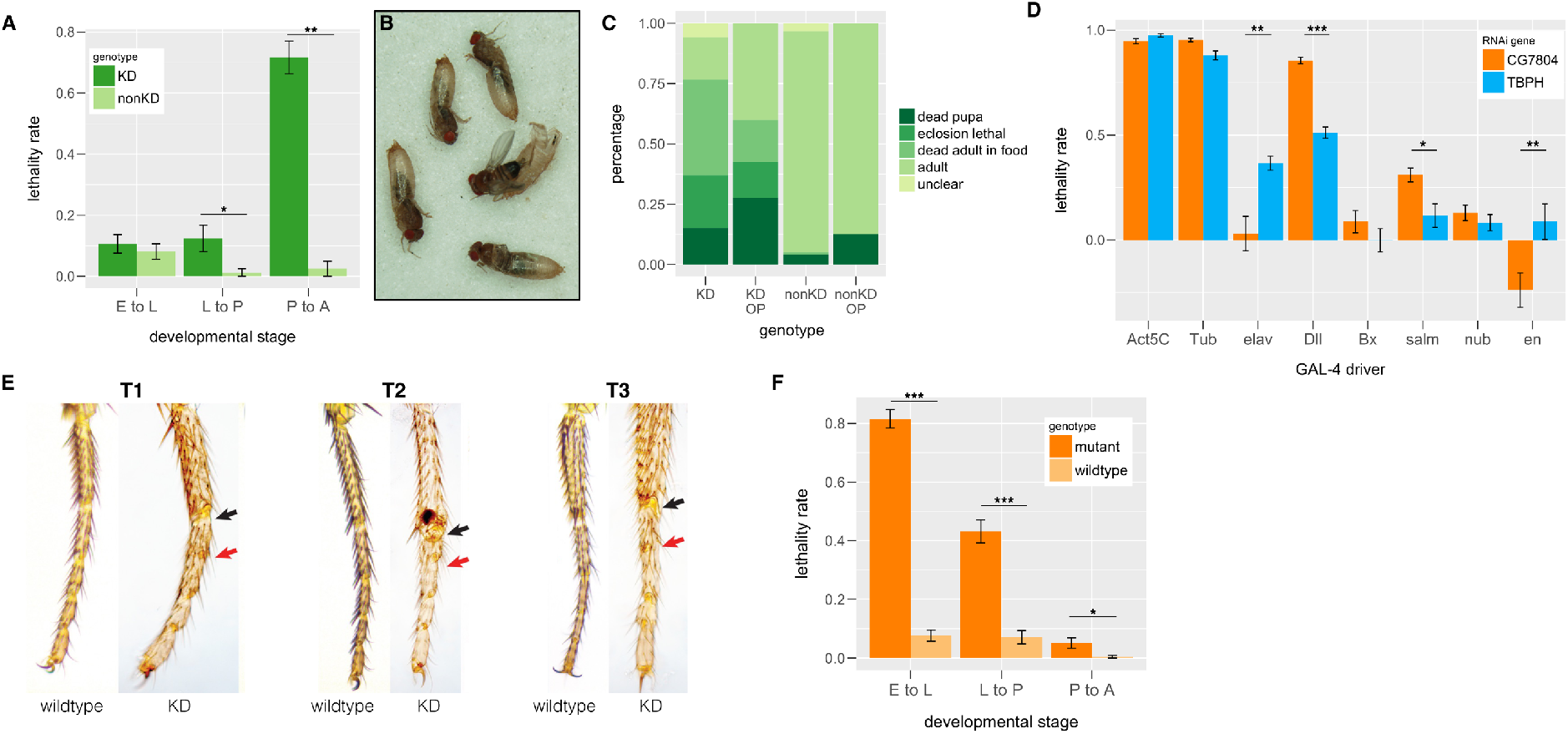
Stage-specific lethality associated with *CG7804* knockdown and knockout. (A) Expression knockdown of *CG7804* using Tub-GAL4 driver results in different lethality rates at different developmental stages. (B) Expression knockdown of *CG7804* leads to eclosion lethal. (C) The outcome of pupae with *CG7804* expression knockdown. (D) *CG7804* expression is essential in different tissues from those of its parental gene, *TBPH.* Lethality rate is significantly different between *CG7804* and *TBPH* knockdown when using *elav, Dll*, and *en GAL4* drivers. Because the lethality rate is estimated relative to wildtype genotype (see Materials and Methods), negative lethality rate (e.g. *en* driver knocking down *CG7804*) means higher survival rate of that particular genotype than the wildtype. (E) Expression knockdown of *CG7804* using Dll-GAL4 driver results in completely fused leg joint (red arrow) or semi-fused leg joint (black arrow). Legs of wildtype (*Dll-GAL4* driver strain) individuals are shown side by side with those of knockdown individuals. T1-T3 are first, second, and third legs respectively. (F) *CG7804* knockout individuals have significantly higher lethality rate from embryo to larva, and from larva to pupa. E: embryo; L: third instar larvae (L3); P: pupae. KD: individuals with *CG7804* knockdown genotype; nonKD: wildtype individuals; OP: pupae with pupa cased removed (open pupae). *Mann-Whitney U test*: \**p* < 0.05, \*\**p* < 0.01, \*\*\**p* < 0.001.

In addition to different developmental stages, we also investigated in which tissues the expression of *CG7804* is essential. According to the modENCODE tissue expression study (Graveley *et al.* 2011; Brown *et al.* 2014), *TBPH* is ubiquitously expressed while *CG7804* is mainly expressed in imaginal discs and male-specific tissues. Because our focus is the evolution of gained essentiality of *CG7804* to survival, we used tissue-specific *GAL4-drivers* to knock down the expression of *CG7804* and *TBPH* in tissues that are not sexually dimorphic. Expression knockdown of *TBPH* using neuronal specific *elva* driver leads to much lower survival rate than expression knockdown of *CG7804* using the same driver (Figure 2E), which is consistent with previously identified role of *TBPH* in neuronal functions (Feiguin *et al.* 2009; Hazelett *et al.* 2012). On the other hand, expression knockdown of *CG7804* using *Dll* (leg imaginal disc) and *salm* (imaginal discs) led to lower survival rate than those of *TBPH* knock down with the same *GAL4 drivers* (Figure 2D). Interestingly, we identified individuals with expression knockdown of *CG7804* at leg imaginal disc (using *Dll GAL4* driver) having fused leg joints (Figure 2E), which is not observed in *TBPH* knockdown flies using the same *GAL4* driver (Supplementary Figure 2). These tissue-specific knockdown results suggest that the expression of *CG7804* is essential for viability at different tissues from those of its parental gene, *TBPH.*

We used CRISPR/CAS9 system (Cong *et al.* 2013; Gratz *et al.* 2013; Kondo and Ueda 2013) to generate null mutant of *CG7804* (see Materials and Methods). Consistent with results using GAL4/RNAi expression knockdown, *CG7804* knock out leads to extremely high lethality rate (99.4%). Interestingly, the lethality associated with *CG7804* knockout happened at earlier stages: mainly at embryo to larva, and larva to pupa stages (Figure 2E), and we did not observe eclosion lethal phenotype with *CG7804* knockout pupa (see Discussion for potential causes). Overall, both our expression knockdown and null mutant analyses support the conclusion that *CG7804*, despite being young and only present in few species, is highly essential for the survival of *D. melanogaster.*

### *CG7804* knock out perturbs expression of genes with important developmental functions

To investigate the mechanisms by which *CG7804* is essential for the survival of *D. melanogaster* at multiple developmental stages, we sequenced and compared the transcriptomes of *CG7804* knockout and wildtype mixed-sex individuals at embryonic, larval, and pupal stages. Comparing between *CG7804* knockout and wildtype individuals, 20.0% (embryo), 6.3% (larva), and 8.5% (pupa) of the genes analyzed have significantly differential expression (False Discovery rate (FDR) < 0.05) (Figure 3A-C). The large number of differentially expressed genes suggests that *CG7804* has a global influence on transcriptome. The especially large number of genes influenced at the embryonic stage is consistent with the observed strong embryonic lethality associated with *CG7804* knockout. At the embryonic stage, the number of down-regulated genes (1,523) in *CG7804* knockout is much greater than that of up-regulated genes (785, binomial test, *p* < 10^-16^, Figure 3A). These down-regulated genes are enriched for Gene ontology of chitin-related processes (e.g. chitin metabolism and catabolism, chitin-based cuticle development (Supplementary Table 1). On the other hand, up-regulated genes are enriched for mitosis-related processes (e.g. chromosome condensation and separation, DNA and centrosome replication, and DNA repair, Supplementary Table 1). In contrast to the observations at the embryonic stage, *CG7804* knock out at larval stage leads to more up-regulated genes, which are surprisingly also enriched with chitin-related processes (653 (up) vs 139 (down), binomial test, *p* < 10^-16^, Figure 3B, Supplementary Table 1). Finally, there are slightly fewer up-than down-regulated genes at the pupal stage with CG7804 knock out (462 (up) vs 585 (down), binomial test, *p* = 0.00016, Figure 3C). Genes with most significant down-regulation again have functions in cuticle development (*Cpr72Eb, Lcp65Ad*). Down-regulated genes are enriched with function in imaginal disc-derived morphogenesis, which is consistent with our findings from tissue-specific expression knockdown analysis (see above, Supplementary Table 1). Chitin is the basis of critical structures of insects (e.g. the exoskeleton and trachea), and insect growth and morphogenesis heavily depend on the synthesis and remodeling of chitin-based structures (Merzendorfer and Zimoch 2003). It is expected that these process will be especially important for the development of embryos (develop into larvae) and pupa (develop into adults), which is consistent with our GO enrichment analyses at these two developmental stages. Intriguingly, why genes with chitin-related functions are instead up-regulated in larvae in *CG7804* knock out is unclear. Interestingly, among all genes analyzed, there seems to be little consistency in the directionality of expressional changes across developmental stages (Figure 3D), suggesting that the global influence of *CG7804* on the transcriptome is contingent on the gene expression network at specific developmental stages.

**Figure 3.**
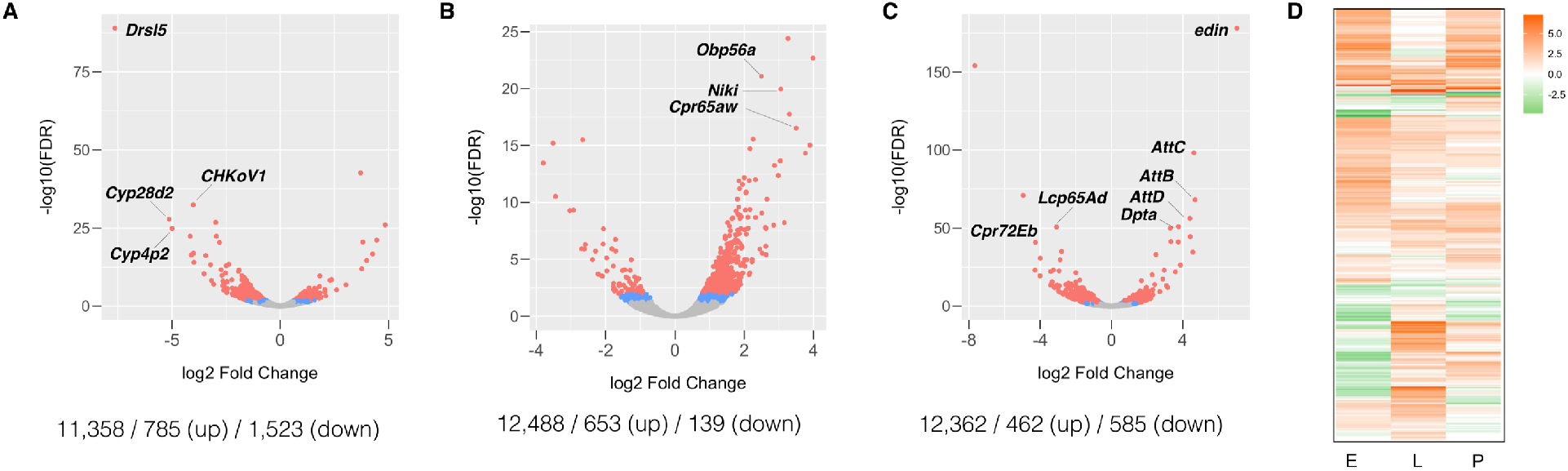
Differential expression upon CG7804 knockout. Volcano plots for the log2 Fold Change in expression level (x-axis) and -log10 FDR (y-axis) for embryonic stage (A), larval stage (B), and pupal stage (C). Red dots represent genes that are differentially expressed with FDR < 0.01 and blue dots for FDR < 0.05. Gray dots represent genes that are not differentially expressed. Numbers under each panel are the number of genes analyzed / number of up-regulated genes / number of down-regulated genes (D) heatmap for the log2 Fold Change in expression level at three developmental stages (E: embryo, L: larva, P: pupa). Each horizontal row represents one gene. Oranges are for positive log2 Fold Change (i.e. up-regulated with *CG7804* knockout) while greens are the opposite.

Previous study found that knock out of *TBPH*, the parental gene, leads to alternative splicing of more than a hundred transcripts (Hazelett *et al.* 2012). However, we did not detect primary transcripts whose alternative spliced form significantly differs between wildtype background and *CG7804* knock out (see Materials and Methods), which could be true biologically or due to not enough sequencing coverage in our dataset to robustly detect differential alternative splicing.

### CG7804 gained novel binding sites

The two predicted RRM domains, which were shown to bind to both DNAs and RNAs in protein TBPH (Kuo *et al.* 2009), harbor substitutions that are unique to *CG7804* and might have significant functional consequences (see above). Accordingly, we hypothesized that *CG7804* quickly became essential through acquiring new nucleic acid interaction partners. Here, we focused on the evolution of potential DNA-binding targets of protein CG7804, and generated transgenic *D. melanogaster* strains that express GFP-tagged CG7804 and GFP-tagged TBPH under endogenous cis-regulatory sequences to test our hypothesis (see Materials and Methods).

The GFP-tagged CG7804 has a nuclear localization, which is similar to that of TBPH and consistent with CG7804’s predicted function of nucleic acid binding (Figure 4A). We performed Chromatin-Immuoprecipitation-sequencing (ChIP-seq) targeting GFP-tagged CG7804 and GFP-tagged TBPH to identify their genomic binding sites at the pupal stage, at which we found high lethality associated with expressional knockdown of *CG7804.* There are 553 genomic regions enriched with CG7804 binding (Irreproducible rate (IDR) < 0.001, see Materials and Methods), and the majority of them overlap with at least one gene (470, 85.0%). This is significant when compared to randomly selected genomic windows with matching chromosomal distributions and sizes (Permutation test, *p* = 0.016). Over a quarter of the remaining enriched region (28, 32.5%) are within 2kb upstream of transcription start sites, also having a potential role in regulating gene expression. In total, we identified 649 genes that either are in CG7804 enriched region (628 genes) or have CG7804-binding enrichment in their upstream 2kb region (358 genes), with 337 genes having CG7804-binding enrichment at both their upstream and gene sequences. A large fraction of genes with CG7804 enrichment also has TBPH enrichment (529, 81.5%). Nevertheless, 120 genes only have CG7804-binding enrichment, which can be regarded as CG7804 gained interaction partners. Interestingly, even for genes that are enriched for the binding of both CG7804 and TBPH, the location of the enrichment region may diverge, which suggests that the paralogs may have differential regulation of some target genes (Figure 4B).

**Figure 4.**
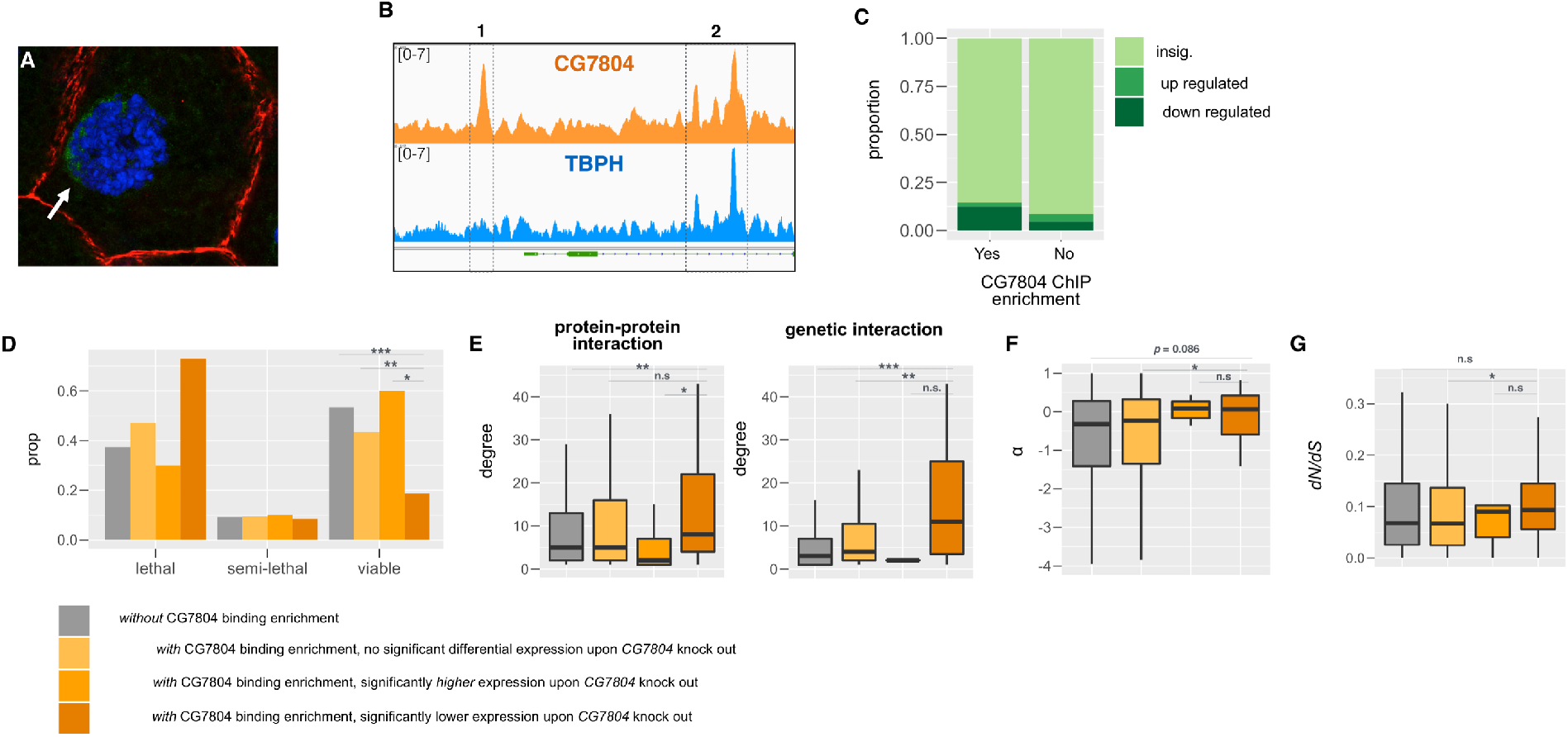
Genes with CG7804-binding enrichment are different from other genes in the genome. (A) CG7804 has nuclear localization in the salivary gland of third instar larva. Green: CG7804, blue: DNA, Red: cytoskeleton. (B) Examples of a binding region that is unique to CG7804 (1) or is shared between CG7804 and TBPH (2). Fold enrichment tracks for CG7804 and TBPH are shown in the first and third tracks respectively. The second and fourth tracks are the identified enrichment region by MACS2 and IDR (see Materials and Methods). This example shows that even for some genes bound by both paralogs, CG7804 may still have gained unique binding sites (here, upstream of an essential gene, MyC). (C) Bar plots for the proportion of genes that are differentially expressed (either down-regulated or up-regulated) for genes with or without CG7804-binding enrichment. Different shade of green colors is whether a gene is differentially expressed upon *CG7804* knockout. (D-G) Comparing (D) known mutant phenotype, (E) degree (number of protein-protein interaction/genetic interaction a gene is involved in), (F) α (the proportion of adaptive amino acid substitution), and (G) *dN/dS* ratio (linage-specific substitution rates) between genes with/without CG7804-binding enrichment and differential expression upon *CG7804* knockout. Three statistical tests are (1) between genes with and without CG7804-binding enrichment, (2) among genes with CG7804-binding enrichment, genes with and without differential expression upon *CG7804* knockout, and (3) among genes with CG7804-binding enrichment and differential expression upon *CG7804* knockout, genes with increased and decreased expression. *Mann-Whitney U test:* *p < 0.05, **p < 0.01, ***p < 0.001.

To investigate if these CG7804-binding enrichment over the genome indeed has functional consequences, in particular influencing the transcript levels of targeted genes, we compared our ChIP-seq results with RNA-seq results of *CG7804* knockouts at the pupal stage (see above). Genes that have CG7804-binding enrichment are also more likely to have significant differential expression (FDR < 0.05, see above) between *CG7804* knockouts and wildtype pupa (*Fisher’s Exact Test, p* < 10^-5^, odds ratio = 1.80, Figure 4C), suggesting a direct effect of the CG7804 binding on expression. In particular, among genes that have CG7804-binding enrichment and are differentially expressed in *CG7804* knockouts, there is an excess of genes that have lowered expression (*Fisher’s Exact Test, p* < 10^-8^, odds ratio = 5.5, Figure 4C). For other genes with CG7804-binding enrichment but not identified as with significant differential expression using our stringent threshold upon *CG7804* knockout (i.e. genes with FDR ≥ 0.05), they still have lower *q-value* (i.e. adjusted *p-value* for controlling for multiple tests, *Mann-Whitney U test p* < 10^-6^) and smaller log2 fold change in expression (i.e. more negative, suggesting lowered expression in knockouts than in wildtype; *Mann-Whitney U test p* = 0.0080). These results suggest that *CG7804’s* role in gene regulation may be predominantly activation of gene expression, which is opposite to TBPH’s role in gene regulation (Hazelett *et al.* 2012).

A potential mechanism by which *CG7804* became essential is through influencing the regulation of other essential genes in *Drosophila.* Consistently, we observed genes with CG7804-binding enrichment are more likely to have known lethal or semi-lethal phenotypes (shown by either knock out mutant or expression knock down, see Materials and Methods) than other genes (*Fisher’s Exact Test, p* < 10^-8^, odds ratio = 1.66, Figure 4D). Furthermore, among genes with CG7804-binding enrichment, genes that are differentially expressed upon *CG7804* knockout are even more likely to have known lethal or semi-lethal phenotype (*Fisher’s Exact Test, p* = 0.0034, odds ratio = 2.35, Figure 4D), and this is mainly driven by genes that have lowered expression upon *CG7804* knockout (comparing between genes with significant increased or decreased transcript levels, *Fisher’s Exact Test, p* = 0.011, odds ratio = 6.31, Figure 4D). It has been repeatedly observed that hub proteins, which have many interaction partners in gene-gene interaction networks, tend to have essential functions (Jeong *et al.* 2001; Yu *et al.* 2004; Batada *et al.* 2007; Blomen *et al.* 2015). Consistent with this view, genes that have CG7804-binding enrichment have more experimentally validated protein-protein interactions or reported genetic interactions than other genes in the genome (degree in PPI network, *Mann-Whitney U test, p* < 0.0016; degree in genetic interaction network, *Mann-Whitney U test, p* < 10^-6^, Figure 4E). In particular, genes that have CG7804-binding enrichment and lowered expression upon *CG7804* knockout have even more genetic interaction partners than other genes with CG7804-binding enrichment but no differential expression (degree in genetic interaction network, *Mann-Whitney U test, p* = 0.0029, Figure 4E). These observations invite the conclusion that *CG7804* became essential mainly through regulating the expression of other essential genes and/or hub genes.

Interestingly, genes with enrichment of CG7804 binding are more likely to be under adaptive evolution (shown by rejection of McDonald-Kreitman test and have an excess of amino acid substitutions than the null expectation) than other genes (*Fisher’s Exact Test, p* = 0.006, odds ratio = 1.54). They also have marginally significantly larger *α* (proportion of amino acid substitutions that are under adaptive evolution, *Mann-Whitney U test, p* = 0.086, Figure 4G), but no faster rates of nonsynonymous substitutions (*dN/dS* ratio, *Mann-Whitney U test, p* = 0.90, Figure 4G). In particular, among genes with CG7804-binding enrichment, those that have differential expression upon *CG7804* knockout have both more adaptive substitutions (α, *Mann-Whitney U test, p* = 0.041, Figure 4F) and faster rates of protein evolution (*dN/dS* ratio, *Mann-Whitney U test, p* = 0.027, Figure 4G) than those that show no differential expression. Our observation revealed that genes with CG7804-binding enrichment, especially those whose expression is expected to be up-regulated by CG7804 binding (i.e. have lowered expression upon *CG7804* knockout), are more likely to have known lethal phenotype, have multiple interaction partners in protein-protein network, and are more often under adaptive evolution.

GO enrichment analysis shows that genes with CG7804-binding enrichment are significantly (*p-value* < 0.05 after multiple test correction) enriched with functions related to development (Supplementary Table 2). This includes imaginal disc-derived morphogenesis, which supports observed lethality of tissue-specific knockdown of *CG7804* in imaginal discs (Fold Enrichment = 4.47), as well as eye, trachea, and neuronal development. Interestingly, genes with CG7804-binding enrichment are also enriched with protein binding (Fold Enrichment = 1.69) and transcription factor activity (Fold Enrichment = 4.34), both of which are expected to influence a large number of genes and have extensive functional impacts. Even more, genes with CG7804-binding enrichment and lowered expression upon *CG7804* knockout show even stronger enrichment for imaginal-disc related morphogenesis (Fold Enrichment = 6.56) and transcription factor activity (Fold Enrichment = 4.65, Supplementary Table 3).

There are 120 genes that only have CG7804-binding enrichment but not TBPH-binding enrichment. The 120 genes that only have CG7804-binding enrichment are not different in terms of their expression, protein/genetic interaction network, and evolutionary rates if compared to those that have binding enrichment for both CG7804 and TBPH. Nevertheless, a large fraction of these genes also shows differential expression upon *CG7804* knockout (19%) and majority of them have down-regulated gene expression (87.5%), which supports the regulatory importance of these CG7804-unique binding enrichment.

## Discussion

The genetic basis of the diversity of life remains a central question in evolutionary biology. Nucleotide substitutions or indels that change protein-coding or regulatory sequences are often observed to contribute to functional, phenotypic, and behavior polymorphism and divergence within and between species (e.g. (Rost *et al.* 2004; Wittkopp *et al.* 2009; Linnen *et al.* 2013; Ding *et al.* 2016), reviewed in (Wray 2007; Barrett and Hoekstra 2011)). However, in addition to gradual change at pre-existing genes, gene composition turns over rapidly even between closely related species (e.g. (Demuth *et al.* 2006; Zhang *et al.* 2010, 2011) and reviewed in (Kaessmann 2010)) Indeed, while humans and chimpanzees have only diverged 1.5% in their orthologous coding sequences (Consortium 2005), they differ by at least 6% of their gene contents (Demuth *et al.* 2006). The origination of new genes is a major contributor to this dynamic gene turn over. Despite being young and restricted to few species on a phylogeny, new genes have diverse essential functions and the rapid evolution of these new genes copies is an important driving force for adaptation (reviewed in (Chen *et al.;* Ventura and Long; Taylor and Raes 2004; Kaessmann 2010; Ding *et al.* 2012; Long *et al.* 2013)). Yet, how, in a short evolutionary time, new genes became essential is still an open question, and detailed functional dissection of new genes is a natural and important step to address this overarching question.

Despite its young age, our focused duplicated gene, *CG7804*, is found essential for *D. melanogaster* viability in both expression knockdown and gene knockout experiments. Interestingly, we identified eclosion lethal (i.e. flies got stuck halfway in pupal cases) as well as fused leg joints phenotype associated with *CG7804* knockdown, and the former led to our naming this gene *“Cocoon”.* The essentiality of *Cocoon* is likely the result of positive selection, which is evidenced by the rapid accumulation of amino acid substitutions (*dN/dS*) from the parental gene since its origination and the large proportion of adaptive amino acid substitutions (α). RNA-seq analysis revealed that *Cocoon* has a widespread effect on transcriptome at multiple developmental stages, consistent with the observed essentiality of the gene through development. In particular, *Cocoon* knockout leads to significant changes in expression of a fifth of genes in embryo, in which the functional consequence is expected to impact both somatic and germline cells, and have a long-lasting effect through development. Importantly, our ChIP-seq analysis at pupal stage suggests that *Cocoon* rapidly became essential by forming interactions with and regulating the expression of multiple pre-existing genes, in particular those that have known lethal phenotypes, have many protein-protein/genetic interaction partners (hub genes), and/or are also experiencing adaptive evolution. To the best of our knowledge, this is the first study that detailed the evolutionary steps for how a very young (less than four million years old) new gene became essential for viability.

According to network theory, “hubs” are central nodes with a larger number of links, and perturbation of hubs is expected to have more widespread influence on the overall network than perturbations of peripheral nodes with few links (for biological networks, reviewed in (Barabási and Oltvai 2004; Barabási *et al.* 2011)). Indeed, in gene-gene interaction network, hub genes are often essential (Jeong *et al.* 2001; Yu *et al.* 2004; Batada *et al.* 2007; Blomen *et al.* 2015). Comparisons of ancient orthologous genes found that the essentiality of genes can evolve via gradual increases in the number of interactions (Kim *et al.* 2012). Similarly, genome-wide analysis in yeast, mouse, and human concluded that, on average, the integration of new genes into pre-existing gene-gene interaction network is a gradual process (Capra *et al.* 2010; Abrusán 2013; Zhang *et al.* 2015). However, a young (less than four million years old) transcription factor with novel fertility functions was found to massively reshape the gene interaction network by preferentially binding to and influencing the expression of genes with sex-biased expression (Chen *et al.* 2012). Similarly, we found that *Cocoon*, a duplicate from another nucleic acid binding protein *TBPH* (Kuo *et al.* 2009), became essential through acquiring many interaction partners in a short evolutionary time. In particular, the binding of COCOON to other essential and/or hub genes may further expedite the evolution of gained essentiality. These observations suggest notable exceptions to the common views that the accumulation of genetic interaction is a slow process, and that new genes play a minimum role in essential cellular functions. This may be especially true for certain classes of genes, such as transcription or splicing factors, which can influence expression of hundreds of genes. New genes with these functions may be more likely to acquire multiple new interactions quickly.

While analyses using gene knockout and expression knockdown both supported a strong role of *Cocoon* in viability, the developmental stages at which *Cocoon* has the strongest influence vary between perturbation methods. Specifically, *Cocoon* knockout results in strongest lethality at the embryonic stage, while *Cocoon* knockdown did not show a viability effect until pupal stage. This could be due to the fact that embryonic development also depends heavily on maternally loaded RNAs, and *Cocoon* was previously categorized as being both maternally deposited and zygotically expressed (Lott *et al.* 2011). On the other hand, incongruence between knockout and knockdown phenotypes has been widely observed in other systems (Souza *et al.* 2006; Daude *et al.* 2012; Kok *et al.* 2015), and a recent detailed study pointed out that genetic compensation in knockout mutant, but not in knockdown individuals, is one of the explanations (Rossi *et al.* 2015), suggesting knockout and knockdown should be viewed as complementing methods.

The large overlap of binding targets between Cocoon and TBPH invite the conclusion that, since its origination, *Cocoon* subfunctionlized the function of *TBPH.* However, several pieces of evidence point towards a more complex evolutionary process. First, the protein sequence and the molecular function of TBPH is highly conserved not only within *Drosophila*, but also across animals (Supplementary Figure 1). In fact, human version of *TBPH* (*TDP-43*) was able to complement loss of function *TBPH* mutant in *D. melanogaster* (Li *et al.* 2010). This would suggest that the function of *TBPH* likely hasn’t changed since the origination of *Cocoon.* Besides, our tissue-specific expression knockdown analysis found that *Cocoon* and *TBPH* support essential functions in different tissues. Importantly, while we identified that COCOON-binding predominantly results in up-regulation of targeted genes, TBPH’s role in expression regulation was more often negative (i.e. down-regulation) in both flies (Hazelett *et al.* 2012) and mice (Polymenidou *et al.*

2011). These observations suggest that *Cocoon* might have evolved new function (neo-functionalized) since its duplication from *TBPH.* Our study provides an unprecedented view of how duplicated new genes became essential for viability rapidly through forming connections with other essential/rapidly evolving genes in the gene interaction network at critical developmental stages.

## Materials and Methods

### Evolutionary genetic analysis of *CG7804* and *TBPH*

We used coding sequence of *CG7804* and *TBPH* of 12 *Drosophila* species from (Clark *et al.* 2007) and aligned using Clustal (Sievers *et al.* 2011), followed by manual curation. We used PAML (Yang 2007) and ran (1) branch model (to estimate number of nonsynonymous and synonymous substitutions on each branch), and (2) clade model (to test for different rates of nonsynonymous substitutions between *CG7804* and *TBPH).* For McDonald-Kreitman test, we used *D. melanogaster* polymorphism data from (Lack *et al.* 2015) and used *D.* simulans allele from (Hu *et al.* 2013) as outgroup to perform unpolarized test.

Domains of CG7804 were predicted by Pfam (Finn *et al.* 2016) and the tertiary structures of predicted domains were computed using Phyre (Kelley *et al.* 2015). To have a broad view of the evolutionary conservation among *TBPH* orthologs, protein sequences from 49 species were retrieved from NCBI, aligned using Clustal (Sievers *et al.* 2011), followed by manual curation. We then compared the residues found at specific functional sites that were tested experimentally by (Lukavsky *et al.* 2013).

### Generation of transgenic strains and mutants

Design of guide RNA, injection of guide RNA, and screen for CRISPR mutants were done by Genetic Service Inc. (Sudbury, MA). *CG7804* mutant has 2 bp deletion in the coding sequence, which is confirmed by Sanger sequencing. Detailed CRISPR design and sequencing confirmation are in Supplementary Text 1. To further confirm that our CRISPR mutant is true null mutant, we used other mutant of *CG7804* to perform complementation tests. We used a strain (BDSC 36014, Supplementary Table 4) that has a MIMIC construct (Venken *et al.* 2011) inserted in the coding sequence of *CG7804*, and is likely a null allele of *CG7804.* The presence of the MIMIC insertion was confirmed by PCR (see, Supplementary Table 5 for primer information). We found CRISPR and the MIMIC strain do not complement each other, suggesting CRISPR *CG7804* is a null mutant of *CG7804* (Supplementary Table 6). *TBPH* mutant is from (Hazelett *et al.* 2012) and we balanced both *CG7804* and *TBPH* mutants over balancer chromosomes with ubiquitously expressed GFP for developmental stage-specific lethality analysis (see Supplementary Table 4 for strains used).

It is worth mentioning that, even for those few *CG7804* knockout individuals that survived to adult (i.e. the lethality rate is not 100%), we were able to detect expression of *CG7804* either through RT-PCR or RNA-seq. However, these detected transcripts of *CG7804* all have the same deletion that causes frame shifts.

Constructs of GFP-tagged *CG7804* and GFP-tagged *TBPH* were generated using BAC-recombineering and P(acman) BACs CH322-116J04 (*CG7804*) and CH321-59A22 (*TBPH*) (Venken *et al.* 2009) and cloning vector pAV007 (GeneBank # KF411445) by Genome Engineering Core of the University of Chicago (Chicago, IL). The pAV007 was inserted directly after CGAACCAGAGCAGCGGATCTCAAAACGCCGCGGAGAAGTCAAACTTTCTT in CH321-59A22 (*TBPH*) and after GCATGCATTCATTTAATCCACATGGTTACCAAATGAATCGCGTCATGAAC in CH322-116J04 (*CG7804*) to generate C-termini GFP-tagged proteins. These constructs were introduced into the genomes of strains with attP docking sites (Bateman *et al.* 2006; Bischof *et al.* 2007)(see Supplementary Table 4 for strains used). Embryo injection of constructs, screening, and balancing were done by Genetivision (Houston, TX). Insertions of BAC constructs were confirmed by PCR (see Supplementary Table 5 for primer sequences).

### Essentiality analysis

Virgin females of RNAi strain (homozygous) were crossed to males of *GAL4* strain. The *GAL4* strain is heterozygous for *GAL4* construct, which is balanced with visible markers and/or construct of ubiquitously expressed GFP. Expression knockdown and wildtype offspring were recognized by visible markers (adult) or presence/absence of GFP (embryo, larva, and pupa). All comparisons are within crosses (RNAi/+; *GAL4/+* vs RNAi/+; GAL4/balancer). For each cross, the expected number of knockdown individuals was estimated using the number of individuals with other genotypes and with the assumption that alleles were inherited following Mendelian rules. The survival rate of knockdown individuals was estimated as observed number of knockdown individuals divided by the expectation, and the lethality rate is one minus the survival rate. At least 10 independent crosses that have at least 20 adults in each cross were counted. For tracking stage-specific lethality, 20 embryos/larvae/pupa of either genotype were collected and placed on fresh medium and the numbers of next-stage individuals were counted after 5 and 10 days. We collected embryos of mixed stages through standard apple juice plate, larvae at L3 stage, and white pre-pupa. This was repeated for at least four times for a specific genotype or specific developmental stage. *GAL4* and RNAi strains used in the study can be found in Supplementary Table 4. Estimation of survival rate and tracking of stage-specific lethality rate for knockout individuals used the same methods. All flies were reared with standard *Drosophila* medium at 25 °C with 12/12 light and dark cycle.

### Real-time RT-PCR analysis

Expression knockdown of *CG7804* and *TBPH* by RNAi was confirmed by real-time RT-PCR analysis (Supplementary Figure 3). RNA was extracted from third instar larvae using Qiagen RNeasy mini kit, digested with DNAase I (Invitrogen) to remove DNAs, and reversed transcribed to cDNA using SuperScript III Reverse Transcriptase (Invitrogen). Real-time RT-PCR was performed using iTaq Universal SYBR Green Supermix (Bio-rad). Quantitative PCR values were normalized using the ΔΔC_T_ method to qRp49 control products.

### RNA-seq experiment and analysis

Knockout individuals were collected by crossing individuals that are heterozygous for the null allele (null/GFP-balancer) and collect F1 without GFP (homozygous for the null allele). For the wildtype counterparts, we used the Cas9 strain from which the knockout mutant was generated (see Supplementary Table 4). We collected 0-24hr embryos using standard apple juice plate, wandering L3, and white pre-pupa. For all three stages, we used mixed-sex individuals. Total RNAs were extracted from collected materials using RNeasy Plus kit (Qiagen). RNA-Seq sequencing library was prepared using Illumina TruSeq and sequenced on Illumina Hi-Seq with 100bp, paired-end reads (IGSB Sequencing core, the University of Chicago).

Raw reads were processed with trim-galore (“Babraham Bioinformatics - Trim Galore!”) to remove adaptors and low-quality bases. Processed reads were mapped to *D. melanogaster* release 6 genome (Hoskins *et al.* 2015) with TopHat (Trapnell *et al.* 2009). Htseq-count (Anders *et al.* 2015) was used to count the number of reads mapping to exons. DESeq2 (Love *et al.* 2014) was used to normalize, and estimate expressional fold enrichment between two *D. melanogaster* genotypes. Only genes with at least 10 mean read counts were included in the analysis. We also used the default independent filtering implemented by DESeq2, which uses normalized counts as filter statistics to filter out genes that have little probability of being significant. To detect differential alternative splicing, we used Cufflinks and Cuffdiffs (Trapnell *et al.* 2010). Cuffdiffs computed relative abundance of different splice variants of a primary transcript and used the square root of Jensen-Shannon divergence as a test statistics to test differential alternative splicing between knock out and wildtype.

### ChlP-seq experiment and analysis

ChIP was performed using modEncode protocol (http://www.modencode.org/) with anti-GFP antibody (from Kevin White’s laboratory). ChIP-Seq sequencing library was prepared using NuGen Ovation Ultralow Library Systems V2 (San Carlos, CA) and sequenced on Illumina Hi-Seq with 100bp, paired-end reads (IGSB Sequencing core, the University of Chicago).

Adaptor sequence and low-quality bases were removed from raw reads using trim-galore. Processed reads were mapped to *D. melanogaster* reference genome release 6 (Hoskins *et al.* 2015) with bwa mem (Li and Durbin 2009). Reads with mapping quality smaller than 30 were removed from the analysis using Samtools (Li 2011). Enrichment (IP with respect to Input) was called using MACS2 (narrow peak mode, (Zhang *et al.* 2008)). We also performed IDR analysis (Li *et al.* 2011) to identify enrichment peaks with lower than 1% irreproducibility rate between replicates. Genes overlapping with enrichment peaks were identified using Bedtools (Quinlan and Hall 2010).

### Analysis of gene properties

Degree of each gene in protein-protein network was estimated as the number of experimentally validated, non-redundant protein-protein interaction using data from BioGrid 3.4 (Stark *et al.* 2006). The degree of each gene in genetic interaction network was estimated as the number of reported genetic interaction on flybase (release 2017, Feb). McDonald Kreitman test (McDonald and Kreitman 1991) and the estimation of *α* (Smith and Eyre-Walker 2002), the proportion of adaptive substitutions, were estimated using sequences of Zambia *D. melanogaster* population (Lack *et al.* 2015) and *D. simulans* as outgroup (Hu *et al.* 2013). Lineage specific *dN/dS* ratio was estimated using PAML (Yang 2007), with *D. melanogaster, D. simulans*, and *D.yakuba* alleles. Phenotypic data for all genes annotated were downloaded from Flybase, which were based on either knock out mutant(s) or expressional knockdown analysis. Genes were classified into three categories: lethal (with at least one lethal phenotype observed), semi-lethal (with no lethal phenotype observed and with at least one semi-lethal phenotype observed), and viable (with no known lethal or semi-lethal phenotype). Genes without phenotypic data were excluded from this analysis. GO enrichment analysis was performed using DAVID with Benjamini-Hochberg correction (Huang *et al.* 2009). Because not all genes are expressed at all developmental stages, GO enrichment analysis for differentially expressed genes used list of genes with high enough expression to be included in our RNA-seq analysis as background list.

### Imaging analysis

Images were acquired with a Zeiss LSM700 confocal using either Plan-Apochromat 20x/0.8 M27 or Plan-Apochromat 40x/ NA 1.3 oil-immersion lens. Salivary glands were dissected in PBS, fixed with 4% formaldehyde in PBS for 15 min and stained with rabbit polyclonal anti-GFP (Torrey Pines, 1:1000) and AlexaFluor-488 conjugated secondary antibodies (1:400, Molecular Probes). Dissected tissues were mounted in SlowFade antifade solution (Invitrogen) after TRITC-phalloidin (Sigma) and DAPI (Molecular Probes) stains.

## Acknowledgement

We thank Jennifer Moran of Genome Engineering core of the University of Chicago, and Alec Victor, Matt Kirkey, Jeffrey Gersch from Kevin White’s lab for hosting YCGL for ChIP experiments. We thank Dr. Mortan for generously sharing *TBPH* null mutant and Dr. White for sharing GFP antibody. Mia Levine, Claus Kemkemer, Benjamin Krinsky, and Nicholas VanKuren provided helpful discussions of the project. We are also grateful to Josie Reinhardt, Maria Vibranovski, and Li Zhao for critically reading the manuscript. YCGL was supported by NIH NRSA F32 GM109676 and Chicago Biomedical Consortium Postdoctoral Research Award PDR-043. IMV was supported by the Science without Borders scholarship (BEX18816/12-6). GRR was supported by NSF Graduate Research Fellowship. ML was supported by NSF1051826 and R01GM116113.

## Supplementary Materials

**Table S1. GO enrichment analysis for differentially expressed genes between wildtype and CG7804 knockout embryo, larvae, and pupae.**

**Table S2. GO enrichment analysis for genes with CG7804-binding enrichment. Table S3. GO enrichment analysis for genes with CG7804-binding enrichment and lowered expression upon *CG7804* knockout.**

**Table S4. Strains used in the study.**

**Table S5. Primers used in the study.**

**Table S6. Complementation tests to confirm CRISPR mutants**. CRISPR mutant of *CG7804* was crossed to the MIMIC mutant of *CG7804.* The numbers of F1 for possible genotypes are reported in this table. The CRISPR mutants cannot complement the corresponding mutants, suggesting it is true knockouts of *CG7804*.

**Supplementary Figure 1.**
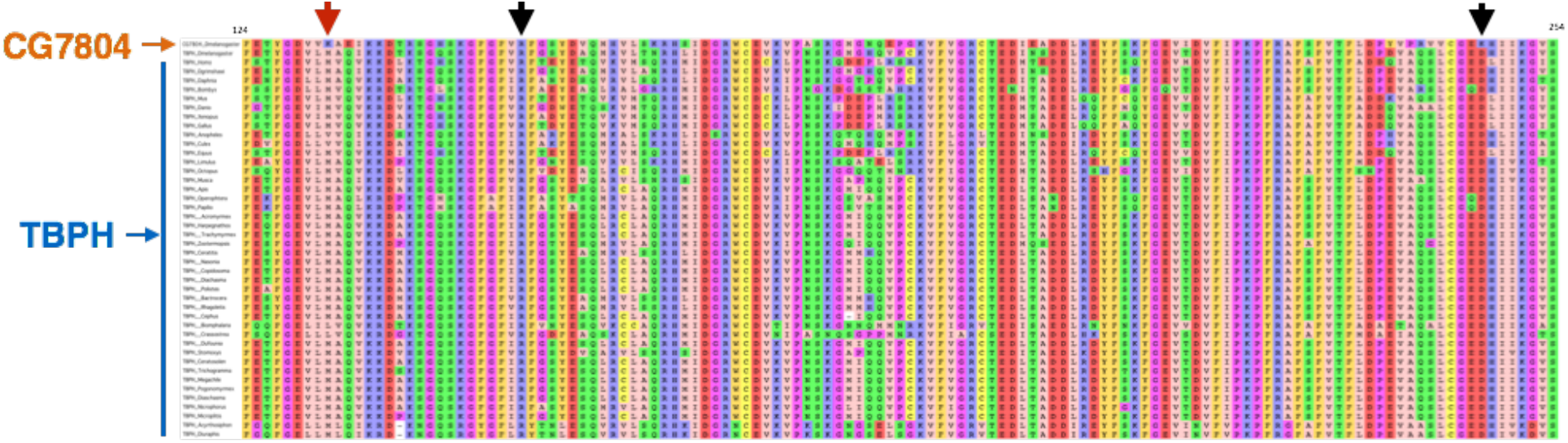
Partial view of the amino acid alignment of TBPH orthologs from 43 animal species and *CG7804* from *D. melanogaster*. The red arrow highlights residue 132, where the methionine was replaced by a lysine in the duplicated protein. Black arrows highlight residues 151 and 247, which interact with each other in the parental protein. Whereas the arginine at position 151 remained conserved, the aspartic acid at residue 247 was replaced for a lysine in the duplicate. These three residues are extensively conserved among TBPH orthologs in animals, and replacements likely have a functional impact in the duplicated protein (Lukavsky *et al.* 2013).

**Supplementary Figure 2.**
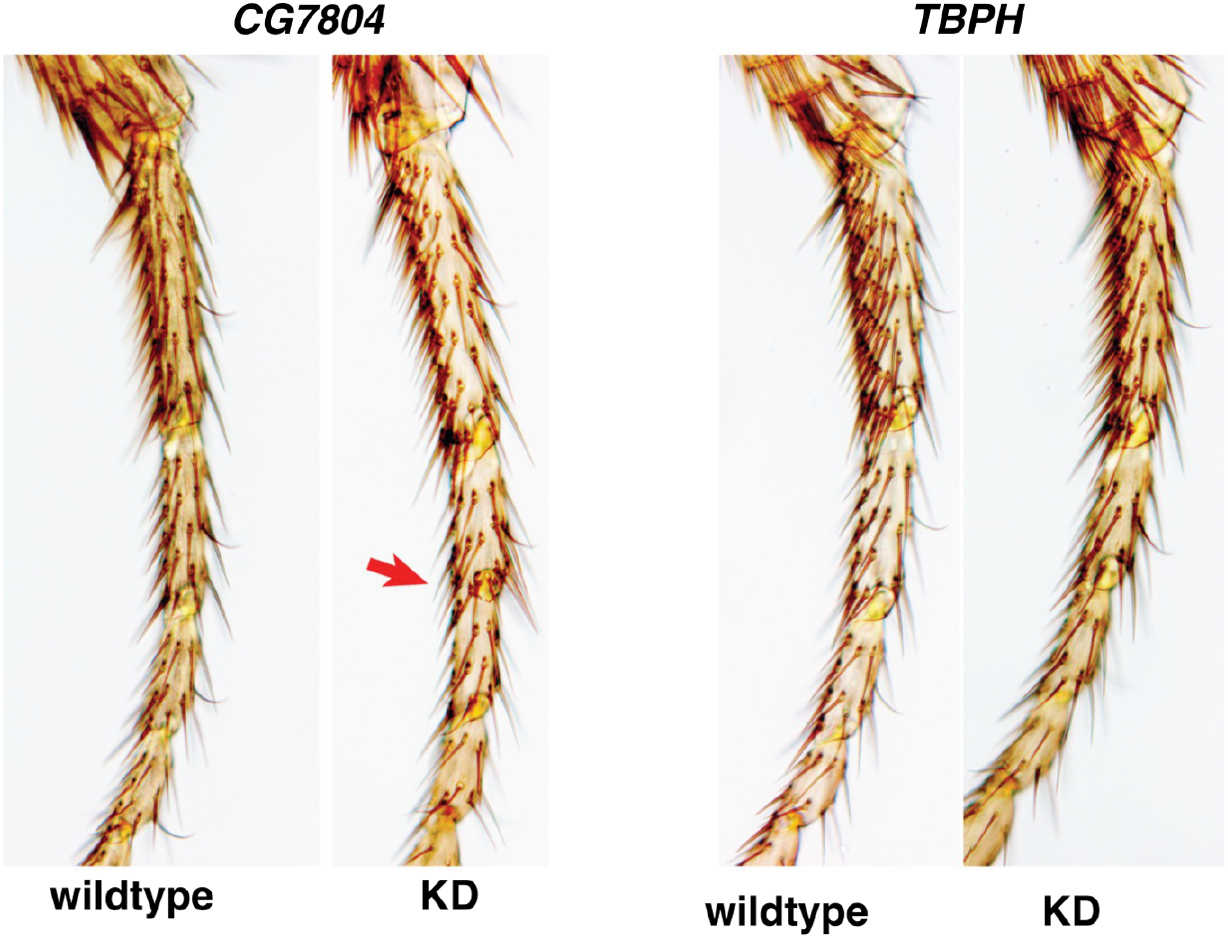
First legs of individuals with *CG7804* or *TBPH* knockdown. Arrows indicate completely fused (red) and semi-fused (black) joints.

**Supplementary Figure 3.**
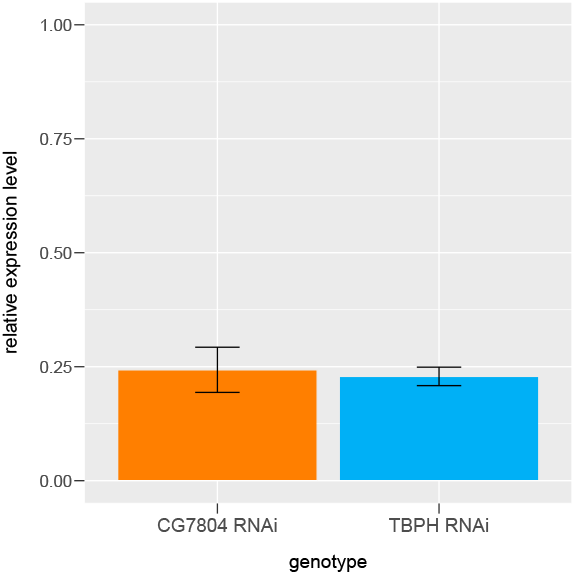
Expression knockdown of *CG7804* and *TBPH* by RNAi. Relative expression levels of *TBPH* and *CG7804* in RNAi knockdowns were compared to wildtype (y-axis). Quantitative PCR values were normalized using the ΔΔCt method, relative to qRp49 control expression. Bars represent means and SEM for three replicates.

**Text S1: CRISPR-guided RNA design and confirmation of CRISPR knockout of CG7804 homozygous individuals**

**CG7804 CRISPR design**

~~~
GTGAGTGCGATCTGAGATCTGGGATCAGAAATCTAAATTTACTCTCTCCATACATAAGACATCAGAATCA GACAGTACAGAACAAATCCCTAGGCAATCCTTATCAATAGTCCGCCGAATAAATGGTTTTCGTTCACGTT TCGGAGAAGGAGGGCGACGAGCCCATCGAGCTGCCGGCTGAGGAGGATGGCACTTTGCTGCTGTCCACAC TGCAGGCGCAGTTTCCGGAATCTAGCGGTCTGCAGTACCGCAACGTGGACACGAAGGCGGTGCGTGGAGT TCGCTCCAACGAGGGCCGACTGTATTCGCCCAGCGAAGAAACCGGCTGGGGCGAGTACCATTACTTTTGC GTCTTTCCCAAGAAGAACAAGCGCCAGAGCGAAGATAACTTGGAAAACTCAACTG**CCAAAA|CCAAGCGC ACTGAGGCC**CATCTGCGCTGCTTTGATCTCATCGTGCTCGGCTTGTCCTATAACACCACCGAGCAGGATT TGCGTGAGTACTTCGAGACCTACGGCGATGTGGTGAAGGCTGAGATCAAGAAGGACACCAGGTCGGGCCA CTCCAAGGGTTTCGGGTTCGTGCGCTTCGGCTCTTACGATGTCCAGATGCACGTACTCTCTAAGCGCCAT TCAATTGATGGTCGCTGGTGCGAGGTTAAGGTGCCCGCCTCGAGGGGCATGGGCAATCAGGAGCCTGGCA AGGTATTCGTCGGCCGCTGCACCGAGGACATAGAGGCGGATGACTTGCGCGAGTACTTCTCAAAGTTTGG CGAAGTGATCGACGTGTTCATTCCCAAGCCCT
~~~

**Bold** indicates CRISPR recognition site

_ (Underscore) labels the CRISPR cut site

**Red** indicates the CRISPR induced deletion

A synthetic guide RNA (bolded text) was designed to direct a Cas9 induced cut between nucleotides 822 and 823 (vertical bar). We generated a mutation with a two base pair deletion directly adjacent to the cut site (red text)

**Sequencing confirmation of CG7804 mutant**

We induced a two base pair mutation of nucleotides 822 and 823. The CRISPR mutant causes a frameshift that results in the a change in amino acids 97-99 and premature stop code inducing a truncation of amino acids 100-318. Red boxes and arrows represent the position of the deleted nucleotides. (A) the alignment of the CG7804 CRISPR line with the genome sequence. (B) cladogram of the CG7804 sequence where nucleotides 822 and 823 have been deleted. (C) alignment of the predicted amino acid sequence of wildtype and CRISPR mutant CG7804 protein.

